# Re-appraising the role of T-cell derived interferon gamma in restriction of *Mycobacterium tuberculosis* in the murine lung

**DOI:** 10.1101/2024.04.04.588086

**Authors:** Karolina Maciag, Courtney Plumlee, Sara Cohen, Benjamin Gern, Kevin Urdahl

## Abstract

T cells producing interferon gamma (IFNγ) have long been considered a stalwart for immune protection against *Mycobacterium tuberculosis* (*Mtb*), but their relative importance to pulmonary immunity has been challenged by murine studies which achieved protection by adoptively transferred *Mtb*-specific IFNγ^-/-^T cells. Using IFNγ^-/-^ T cell chimeric mice and adoptive transfer of IFNγ^-/-^T cells into TCRβ^-/-^δ^-/-^mice, we demonstrate that control of lung *Mtb* burden is in fact dependent on T cell-derived IFNγ, and furthermore, mice selectively deficient in T cell-derived IFNγ develop exacerbated disease compared to T cell-deficient controls despite equivalent lung bacterial burdens. Deficiency in T cell-derived IFNγ skews infected and bystander monocyte-derived macrophages (MDMs) to an alternative M2 phenotype, and promotes neutrophil and eosinophil influx. Our studies support an important role for T cell-derived IFNγ in pulmonary immunity against TB.

## INTRODUCTION

Studies in mice and in human cells have repeatedly demonstrated that immunity to *Mtb* requires both T cells and IFNγ. Genetic deficiencies in CD4+ T cells are associated with increased susceptibility to *Mtb* in mice(1) and humans(2), an outcome mirrored during acquired CD4+ deficiency in advanced HIV(3). Global knockout studies have likewise confirmed the importance of IFNγ in *Mtb* restriction in mice(4, 5), and deficits in IFNγ signaling are associated with human susceptibility to mycobacterial infections – most often with environmentally ubiquitous non-tuberculous mycobacteria, but also TB (6). Nevertheless, the relative contribution of T cell-derived IFNg to protective immunity remains unclear. Currently, vaccine candidates are routinely assessed on their ability to elicit T cell memory responses – particularly, the ability of peripheral blood T cells to produce interferon gamma (IFNγ) - but this parameter does not consistently correlate with protective immunity to *Mtb*(7–14). Additionally, IFNγ-producing T cells can contribute to amelioration(15) or exacerbation(16) of lung pathology in the context of *Mtb* infection. Studies on the importance of polyfunctional CD4+ T cells, as well as in vivo(15–18) and in vitro(19) studies suggestive of IFNγ-independent CD4+ T cell functions, have cast doubt on the primacy of T cell-derived IFNγ. IFNγ is produced by other cell types in addition to T cells during *Mtb* infection(20), and the relationship between IFNγ and its source and target cells remains incompletely understood. Understanding the relative role of T cell-derived IFNγ will inform a rational approach to inducing vaccine-mediated protection against tuberculosis(11, 14).

Prior T cell adoptive transfer studies in mice on the C57BL/6 genetic background have challenged the relative importance of T cell-derived IFNγ in immunity to *Mtb*. In *Mtb*-infected RAG2^-/-^ host mice, adoptive transfer of CD4+ T cells from previously *Mtb*-infected, antibiotic-treated donor mice reduced *Mtb* lung burden and conferred a survival advantage relative to no-transfer control mice whether or not donor T cells expressed IFNγ(10). In another study, adoptive transfer of naïve CD4+ T cells from uninfected donor mice into *Mtb*-infected RAG1^-/-^ mice reduced *Mtb* burden in the lung in a manner only partially dependent on donor T cell IFNγ(16). Finally, in WT mice, adoptive transfer of *in vitro* Th1-polarized CD4+ TCR-transgenic T cells specific for the immunodominant *Mtb* antigen ESAT6 reduced *Mtb* burden in a manner partially – or, at high doses of transferred cells, completely – independently of donor T cell-derived IFNγ(18). Together, these reports suggest that CD4+ T cell-derived IFNγ plays a minimal(18) or partial(15, 16) role in protective immunity against *Mtb* within the lung.

While these studies highlight potential roles of T cell immunity beyond IFNγ, the adoptive transfer techniques employed carry limitations that may lead to underestimation of the relative importance of T cell-derived IFNγ. For example, transfer of a fixed quantity of T cells into lymphopenic mice leads to a limited T cell repertoire (21), potentially skewing the resulting immune response. In this scenario, adoptively transferred T cells also undergo rapid proliferation, differentiation, and activation(21, 22), which would alter their response to subsequent infection and can lead to systemic autoimmunity(23). In addition, RAG-deficient host mice lack not only T cell function but also lack mature B cells and normal lymphoid structure(24). In TCR-transgenic adoptive transfer studies, a very large number of monoclonal *Mtb*-specific T cells may amplify the relative contribution of IFNγ-independent mechanisms of protection that may play a minor role in a more physiologic *Mtb*-specific cell response. Here, we use a T cell bone marrow chimera model to address some of the shortcomings of prior adoptive transfer studies and to re-assess the hypothesis that T cell-derived IFNγ plays only a minor role in T cell-dependent immunity to *Mtb* in the lung. In contrast to methods used in prior studies, we use the more specific T cell-deficient TCRβ^-/-^δ^-/-^host strain, and the T cell chimera model allows physiologically relevant T cell development, including generation of a diverse TCR repertoire, thymic selection, and homeostatic regulation, to occur in the host mouse (25). Our findings indicate that T cell-derived IFNγ is indeed essential for pulmonary immune protection against *Mtb*, providing a reappraisal of the relative importance of this aspect of T cell mediated immunity.

## MATERIALS AND METHODS

### Mice

TCRβ^-/-^δ^-/-^ (strain #002122), IFNγ^-/-^ (strain #002287), and C57BL/6J (wildtype, strain #000664) mice were obtained from Jackson Laboratories (Bar Harbor, ME). Mice were matched by age (when possible) and sex. Mice across different experimental groups were co-housed in the same cage in order to minimize confounding differences in environment and microbiota. Mice were sacrificed using cervical dislocation. All animals were housed and maintained in specific-pathogen-free conditions at Seattle Children’s Research Institute (SCRI). All animal studies were performed with approval of the SCRI Animal Care and Use Committee.

### Bone marrow chimeras

Recipient TCRβ^-/-^δ^-/-^mice were sublethally irradiated with 600 rads. Bone marrow was prepared from femurs and tibias of TCRβ^-/-^δ^-/-^, WT, and IFNγ^-/-^ donor mice, and depleted of mature T cells using a CD3ε depletion kit (Miltenyi Biotec, #130-094-973). 1-2×10^6^ bone marrow cells were then injected retro-orbitally to each host mouse under isoflurane anesthesia. Mice received enrofloxacin in drinking water for four weeks after irradiation to prevent neutropenic sepsis. Chimeric mice were rested for 8-10 weeks prior to subsequent manipulation to allow for T cell development.

### Adoptive transfer

Adoptive transfer of total CD4 T cells into immunocompromised host mice was performed 7 days after aerosol *Mtb* infection of host TCRβ^-/-^δ^-/-^ or RAG1^-/-^ mice as follows. CD4+ T cells were isolated from spleens and lymph nodes of donor C57BL/6 and IFNγ^-/-^ mice using the MagniSort CD4+ T cell enrichment kit (ThermoFisher, #8804-6821-74). Flow cytometry confirmed that >85% of isolated live CD4+ T cells were naïve (CD44loCD62Lhi). 3×10^6^ CD4+ T cells were injected retro-orbitally to each host mouse under isoflurane anesthesia.

### Mtb aerosol infections

T cell chimeric mice were infected with 25-100 CFU aerosolized *Mtb* H37Rv transformed with a reported plasmid bearing the mCherry fluorescent marker constitutively expressed under the pMSP12 promoter sequence(26). Adoptive transfer mice were infected with 25-100 CFU aerosolized *Mtb* H37Rv. Aerosol infections were performed in a Glas-Col chamber. Two additional mice in each infection were sacrificed directly after infection to confirm the infectious dose of *Mtb* CFU per mouse.

### CFU plating

Mouse lung lobes and spleens were homogenized in M tubes (Miltenyi Biotec) containing 1mL PBS+0.05% Tween-80 (PBS-T) using a GentleMACS tissue dissociator (Miltenyi Biotec). Organ homogenates were then diluted in PBS-T and aliquots were plated onto 7H10 agar media to quantify *Mtb* burden. Plates were incubated at for 37°C for at least 21 days prior to CFU counting.

### Flow cytometry

CD45.2 antibody (0.2 mg, PE) was injected into mice retro-orbitally 5-10 minutes prior to sacrifice to label intravascular cells. Lung lobes were excised and dissociated in C tubes (Miltenyi Biotec) in HEPES buffer containing Liberase Blendzyme 3 (70 mg/ml; Roche) and DNaseI (30 mg/ml; Sigma-Aldrich) using a GentleMACS dissociator (Miltenyi Biotec). Lung homogenates were incubated for 30 minutes at 37°C and further processed with the GentleMACS dissociator. Cell suspensions were filtered through a 100 μm cell strainer, treated with RBC lysis buffer (Thermo), and resuspended in FACS buffer (PBS containing 2.5% FBS and 0.1% NaN3). Single cell suspensions were washed in PBS and then incubated with 50 μl Zombie Aqua viability dye (BioLegend) for 10 minutes at room temperature in the dark. Cell markers were stained and viability dye was quenched by the addition of 100 μl of a cocktail of fluorophore-conjugated antibodies diluted in 50% FACS buffer/50% 24G2 Fc block (Bio X Cell, 2.4G2), and incubated for 20 minutes at 4°C. Cells were washed once with FACS buffer and fixed with 1% paraformaldehyde for 30 minutes prior to analysis on an LSRII flow cytometer (BD Biosciences).

### Cell sorting

Lungs were processed as described for flow cytometry above, but NaN3 was omitted from FACS buffer. Cells were sorted on a FACSAria cell sorter (BD Biosciences) under BSL3 conditions.

### RNA-seq

Single cell suspensions from lung were analyzed and sorted by fluorescence-activated flow cytometry (FACS) into *Mtb*-infected (mCherry+) and bystander (mCherry-) MDM (dead-SiglecF-Ly6G-CD11b+ CD64+) populations. RNA was isolated using Trizol, and quantified using bulk RNA-seq (Psomagen) after construction of Illumina sequencing libraries using the SMARTer Stranded Total RNA-Seq Kit v3 - Pico Input Mammalian (Takara). Noise from low-expression transcripts was filtered, and analysis of differentially expressed genes (DEGs) across groups was done using the edgeR module in R(27).

### Protein quantification

Lung lobes were homogenized in M tubes (Miltenyi Biotec) containing 1ml of ProcartaPlex Cell Lysis Buffer (Invitrogen EPX-99999-000) with Halt Protease Inhibitor (Invitrogen 78440) and DNase (30 mg/ml; Sigma-Aldrich) using a GentleMACS tissue dissociator (Miltenyi Biotec). Homogenates were centrifuged to pellet debris, and supernatants were filtered twice through a 0.22 μm pore size Costar SpinX column (Corning) to exclude mycobacteria, frozen at −80°C, and assayed after a single freeze-thaw cycle. Total protein was measured by bicinchoninic acid (BCA) assay (ThermoFisher), and these values were used to normalize individual analyte levels in each sample. IL-4, IL-5, and IL-13 levels in lung homogenates were measured using Cytokine Bead Array Flex Sets (BD); bead fluorescence was measured on an LSRII flow cytometer (BD Biosciences) and analyzed by four-parameter log-logistic curve-fitting to the standard curve. IFNγ levels were quantified using a magnetic Luminex assay (ThermoFisher Scientific) and analyzed using BioPlex Manager software (Bio-Rad).

### Histopathologic analysis and confocal imaging

Right inferior lung lobes were dissected and fixed in 20ml of 1:3 dilution of BD Cytofix Buffer (∼1% formaldehyde) for 24hr at 4°C to ensure killing of *M. tuberculosis*, equilibrated in 30% sucrose solution for another 24hr at 4°C, then rapidly frozen in OCT in an ethanol-dry ice slurry and stored at −80°C. For histopathologic analysis, tissue was embedded in paraffin, 4mm tissue sections were prepared with a cryostat and mounted on glass slides, stained with hematoxylin-eosin by the UW Comparative Pathology Core Facility, then assessed by a trained veterinary pathologist blinded to group assignments. For confocal imaging, 20mm tissue sections were prepared with a cryostat and mounted on glass slides.

Sections were stained with fluorophore-conjugated antibodies and Nucspot 750/780 nuclear stain (Biotium) overnight at room temperature and coverslipped with Fluoromount G mounting media (Southern Biotec). Images were acquired on a Leica Stellaris 8 confocal microscope, compensated for fluorophore spillover using LAS X (Leica), and rendered in Imaris (Bitplane), where ARG1 signal was smoothed using a Gaussian filter with a width of 0.316μm. Identical settings were applied across experimental groups.

### Statistical analysis

Statistical significance was determined using the multcomp and rstatix packages in R, using methods indicated in the figure legends. Principal component analysis was done using the stats package in R.

## RESULTS

### T cell-derived IFN***γ*** is required to reduce lung Mtb burden, and protect from disease

To investigate the role of T cell-derived IFNγ in TB immunity, we generated T cell chimeric mice (**Fig. 1A**) in which T cells, but not other cell types, were genetically deficient in their capacity to express IFNγ. To establish this system, TCRβ^-/-^δ^-/-^host mice were partially myeloablated by sublethal irradiation and reconstituted with bone marrow of IFNγ^-/-^ donor mice, or as controls, bone marrow of TCRβ^-/-^δ^-/-^ or WT mice. In these chimeras, all T cells are derived from the donor bone marrow (e.g., IFNγ^-/-^ for the experimental group), whereas >95% of other hematopoietic lineage cells remain wildtype due to the sublethal dose of radiation(25). After immune reconstitution, T cell chimeric mice were infected with aerosolized *Mtb* H37Rv, then assessed for bacterial burden in lungs and spleens at 25 days post infection (dpi). In contrast to prior T cell transfer studies suggesting that T cell-derived IFNγ may be partially or wholly dispensable for protective T cell responses against murine pulmonary *Mtb*(16, 18), we found that control of bacterial burden in lungs and spleens of T cell chimeric mice at 25dpi was indeed dependent on T cell-derived IFNγ (**Fig. 1B-C**). Furthermore, IFNγ^-/-^T cell chimeric mice exhibited clinical deterioration (decreased activity, hunched posture) and lost weight beyond 25dpi, while TCRβ^-/-^δ^-/-^ chimeric controls did not (**Fig. 1D**), despite equivalent lung bacterial burdens, suggesting that T cell activity during *Mtb* infection promotes disease unless countered by T cell-derived IFNγ.

**Figure 1:**
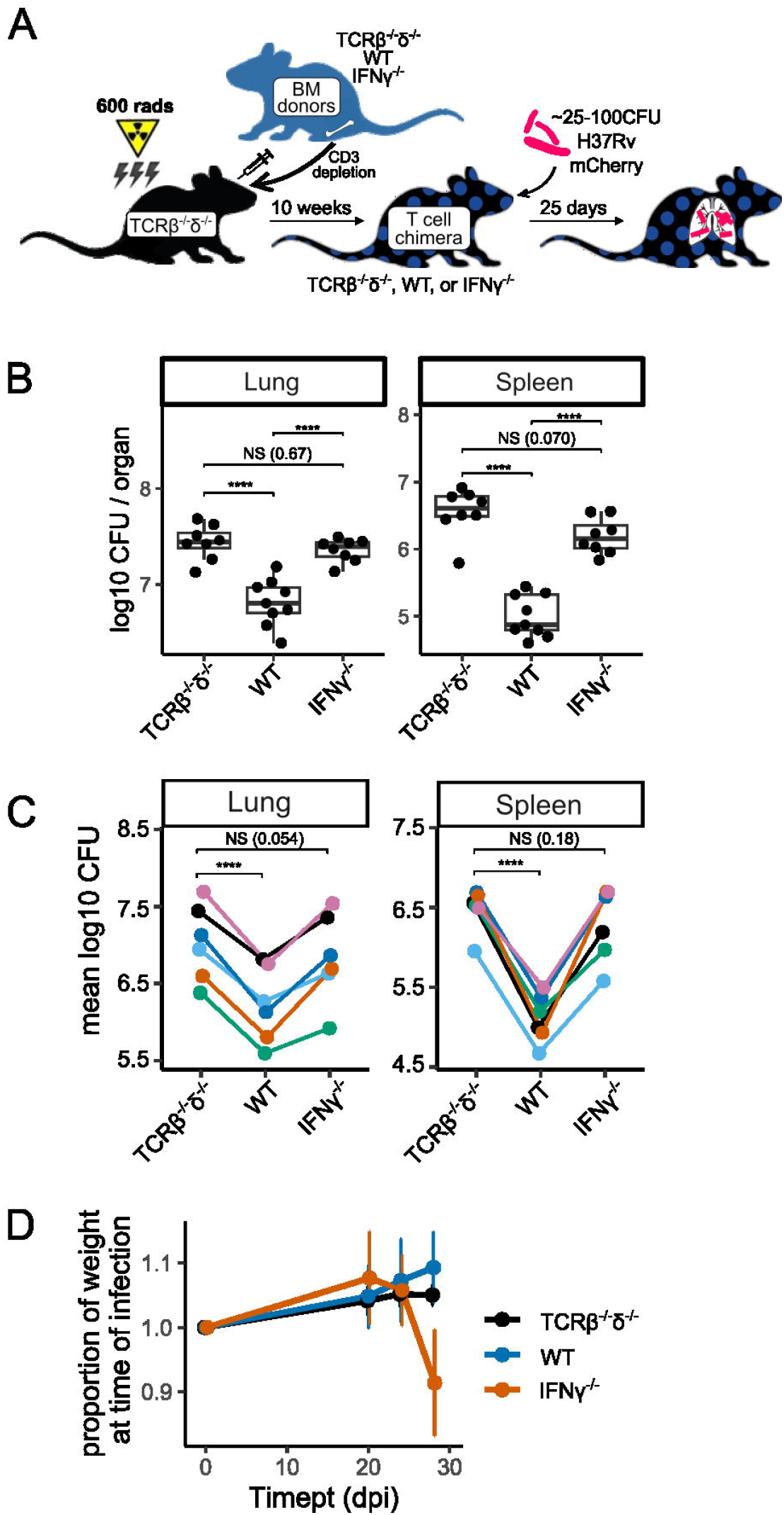
IFNγ^-/-^ T cells do not reduce *Mtb* bacterial burden, and exacerbate disease in T cell chimeric mice. (A) Schematic of the preparation of TCRβ^-/-^δ^-/-^, WT, and IFNγ^-/-^, T cell chimeric mice, followed by infection with aerosolized *Mtb*. (B-C) Bacterial burden in lungs and spleens of *Mtb-*infected T cell chimeric mice at 25dpi in one representative experiment (B), as well as group means from six independent experiments (C). Total n=35 TCRβ^-/-^δ^-/-^, n=34 WT, and n=33 IFNγ^-/-^ T cell chimeric mice. Statistical significance was determined by Tukey’s range test. \**p* ≤ 0.05, \*\**p* ≤ 0.01, \*\*\**p* ≤ 0.001, \*\*\*\**p* ≤ 0.0001. (D) Weight trends of *Mtb-*infected T cell chimeric mice through 29dpi.

In contrast to our results in the T cell chimera model, prior studies have shown that adoptive transfer of IFNγ^-/-^ T cells into T cell-deficient (RAG1^-/-^) host mice decreases lung and spleen *Mtb* CFU relative to no transfer, though the protective effect was smaller than that observed after adoptive transfer of WT T cells(16). We used a similar adoptive transfer strategy (**Fig. 2A**) to reconcile those findings with our results. Instead of RAG-deficient host mice as in Sakai et al., however, we used TCRβ^-/-^δ^-/-^ host mice (as we had used in the T cell chimera experiments, **Fig. 1A**). TCRβ^-/-^δ^-/-^ host mice were infected with aerosolized *Mtb* and CD4+ T cells (3×10^6^/mouse) isolated from WT or IFNγ^-/-^ donor mice were administered intravenously at 7dpi. In order to investigate whether IFNγ^-/-^ T cells mediate pathologic effects, and whether WT T cells can counteract those effects in this model, an additional group of mice received a 50% / 50% mix of WT and IFNγ^-/-^ T cells. Similarly to our observations in T cell chimeric animals, WT T cells significantly decreased *Mtb* burden in both the lungs and spleens of TCRβ^-/-^δ^-/-^ host mice at 36dpi, while IFNγ^-/-^ T cells did not (**Fig. 2B**), again supporting a requirement for T cell-derived IFNγ for control of bacterial burden in murine *Mtb* infection. Furthermore, mice receiving adoptively transferred IFNγ^-/-^ CD4+ T cells exhibited significantly more weight loss over the course of infection than either no-transfer controls or WT CD4+ T cell transfer mice (**Fig. 2C**), despite having equivalent mycobacterial burdens as no-transfer controls. This is again consistent with a pathologic effect mediated by T cells that are unable to produce IFNγ, as observed in T cell chimeric mice. Interestingly, co-administration of WT and IFNγ^-/-^donor CD4+ T cells restored the ability of recipient mice to control mycobacterial burden in lungs and spleens (**Fig. 2B**, “mix”), and decreased the rate of weight loss after infection compared to mice receiving IFNγ^-/-^ T cells alone. This suggests that IFNγ^-/-^ deficient T cells are not inherently pathogenic, and can be complemented *in trans* by the presence of other T cells that can produce IFNγ – but not by other IFNγ-producing cell types (such as NK or NKT cells) during *Mtb* infection.

**Figure 2:**
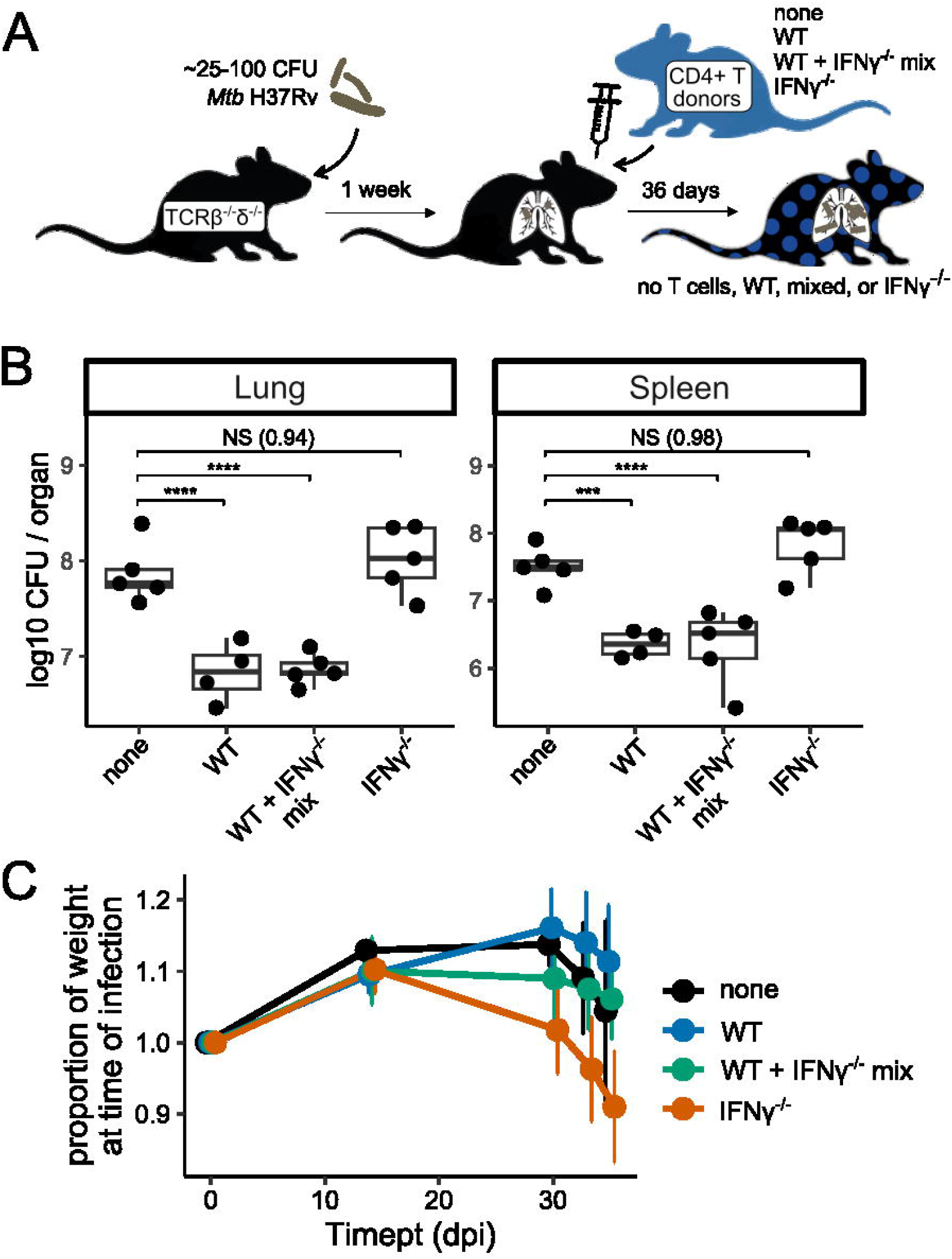
Adoptively transferred IFNγ^-/-^ CD4+ T cells do not reduce *Mtb* burden, and exacerbate disease in TCRβ^-/-^δ^-/-^mice. (A) Schematic of adoptive transfer of zero (none) or 3*10^6^ WT, 50/50% mixed, or IFNγ^-/-^ CD4+ T cells to T cell-deficient host mice after infection with aerosolized *Mtb*. (B) Bacterial burden in lungs and spleens of *Mtb*-infected adoptive transfer mice at 36dpi. Results are representative of two independent experiments. Statistical significance was determined by Tukey’s range test. \**p* ≤ 0.05, \*\**p* ≤ 0.01, \*\*\**p* ≤ 0.001, \*\*\*\**p* ≤ 0.0001. (C) Weight trends of *Mtb*-infected adoptive transfer mice through 36dpi.

In a parallel experiment, we also tested whether the difference between our results and those reported by Sakai et al. could be explained by differences in the host mice used, as ours were specifically deficient in T cells (TCRβ^-/-^δ^-/-^), whereas Sakai et al. used mice deficient in both B and T cells (RAG1^-/-^) (**Fig. S1A**). While Sakai et al. reported 60-fold and 5-fold decreases in lung and spleen *Mtb* burden in RAG1^-/-^ mice that had received IFNγ^-/-^ T cells compared to no-transfer control mice at 42dpi(16), we observed similar lung and spleen mycobacterial burdens in these groups (**Fig. S1B**). However, we had to assess mycobacterial burden in RAG1^-/-^ host mice at the earlier 34dpi timepoint since in our laboratory, *Mtb*-infected RAG1^-/-^ host mice started to lose weight after 23 days of infection regardless of transferred T cell genotype (**Fig. S1C**) and, in a separate experiment, many in fact required euthanasia prior to 42dpi. In addition, within-group variance among lung *Mtb* CFU in RAG1^-/-^ host mice in our hands was high, which limited our ability to observe statistically significant intra-group effects in lung *Mtb* burden (**Fig. S1B, left**) and thus may have also contributed to discordance between our findings and those of Sakai et al. Although our results do not fully explain the discrepancy between our findings and those previously published, they suggest that using RAG1^-/-^ recipients that lack both T cells and B cells and that have aberrant lymph nodes may lead to confounding factors that increase variability in some settings. Taken together, our data using both T cell chimeric mice and adoptive transfer into T cell-deficient mice suggest that T cell derived IFNγ is required for pulmonary immunity against *Mtb*, and that a T cell-specific incapability to produce IFNγ can in fact promote detrimental pathologic effects.

### TB lesions in IFN***γ***^-/-^ T cell chimeric mice exhibit increased neutrophilic and eosinophilic infiltration

To investigate the immune landscape associated in mice with T cell intrinsic IFNγ deficiency, we analyzed the cellular composition of lung tissue of T cell chimeric mice at 25dpi using flow cytometry (**Fig. 3A**, gating as in **Fig. S2A**). Strikingly, the number of both neutrophils and eosinophils in IFNγ^-/-^ T cell chimeric mice was approximately ∼1 log higher than in either TCRβ^-/-^δ^-/-^ (lacking T cells) or WT T cell chimeras (**Fig. 3A**), though T cell-dependent recruitment of monocyte-derived macrophages (MDMs) was preserved in both IFNγ^-/-^ and WT T cell chimeras. Consistent with these findings, confocal microscopy revealed robust neutrophil and eosinophil infiltration into pulmonary TB lesions in IFNγ^-/-^ chimeric mice (**Fig. 3B**).

**Figure 3:**
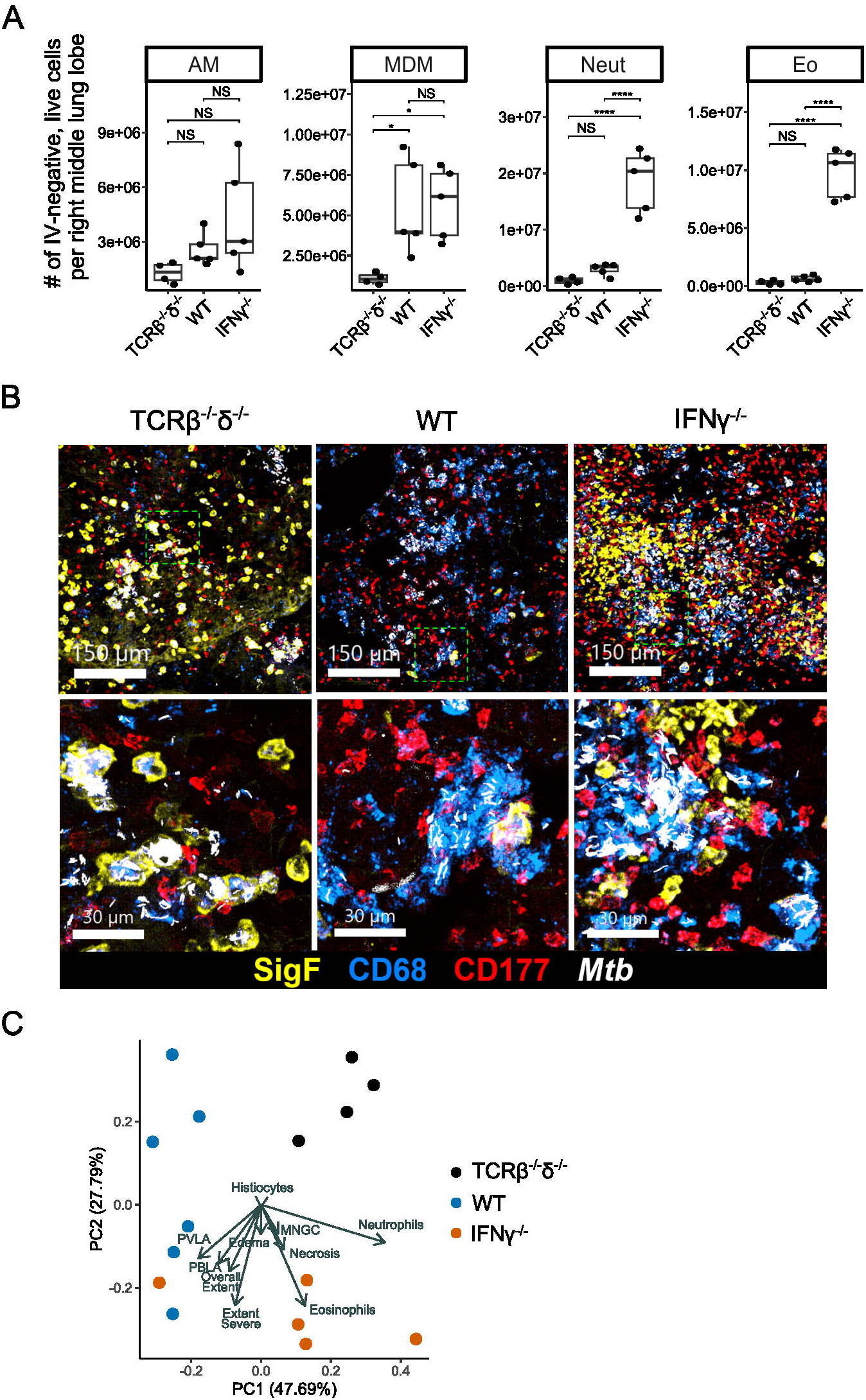
IFNγ^-/-^ T cells promote neutrophil and eosinophil recruitment to pulmonary lesions in T cell chimeric mice infected with *Mtb*. (A) Absolute number of each indicated cell type among live, parenchymal (IV-) cells in the right lung of T cell chimeric mice at 25dpi. Results are representative of six independent experiments. Statistical significance was determined by Tukey’s range test. \**p* ≤ 0.05, \*\**p* ≤ 0.01, \*\*\**p* ≤ 0.001, \*\*\*\**p* ≤ 0.0001. AM: Alveolar macrophage, MDM: monocyte-derived macrophage, Neut: neutrophil, Eo: eosinophil (B) Representative confocal microscopy demonstrating AMs (SiglecF+, CD68+), MDMs (SiglecF-, CD68+), neutrophils (CD177+), and eosinophils (SiglecF+, CD11c-) in TB lesions in T cell chimeric mice. (C) Representative images of TB lesions in H&E-stained sections of T cell chimeric mice at 25dpi. 40x: arrows represent TB lesions. 200x: asterisks represent perivascular and peribronchiolar lymphocyte aggregates. 400x: arrows represent neutrophilic infiltrates. (D) Principal component analysis of fifteen histopathologic features assessed in representative sections of fixed and hematoxylin-eosin (H&E) stained lung of T cell chimeric mice at 25dpi. PVLA: perivascular lymphoid aggregates. PBLA: peribronchial lymphoid aggregates. MNGC: multinucleated giant cells.

We next asked whether histopathologic tissue analysis might help give insight into how increased neutrophil and eosinophil responses (**Fig. 3A-B**) may be linked with clinical decline in IFNγ^-/-^ T cell chimeric mice relative to TCRβ^-/-^δ^-/-^ and WT T cell chimeras (**Fig. 1D**). Hematoxylin-eosin stained lung sections were scored in a blinded fashion across eleven standardized histopathologic features (**Fig. S3A**) confirming that *Mtb* lesions in IFNγ^-/-^ T cell chimeric mouse lungs were marked by abundant neutrophil and eosinophil infiltration (**Fig. S3B**, 800x). Absence of T cells correlated with fewer and more poorly organized TB lesions in TCRβ^-/-^δ^-/-^ T cell chimeric mice, while both IFNγ^-/-^ and WT T cell chimeric mice were able to form dense, organized TB lesions (**Fig. S3B**, 25x). While the differences in histopathology across groups were individually subtle, combined assessment using principal component analysis of the blinded histopathology feature scores clustered samples within each genotype together, indicating similar pathology; this was driven mainly by lesion neutrophils and eosinophils in IFNγ^-/-^ T cell chimeric mice, and decreased extent and severity of lung involvement, as well as absence of lymphoid aggregates, in TCRβ^-/-^δ^-/-^ chimeric mice (**Fig. 3C**). In summary, T cells were able to promote organized TB lesions in lung tissue of T cell chimeric mice during *Mtb* infection regardless of their ability to produce IFNγ, but T cell-derived IFNγ was required to restrict neutrophil and eosinophil infiltration of these lesions.

### IFN***γ***^-/-^ T cell chimeric mice exhibit a Th2 cytokine milieu and alternative activation of MDMs

To further investigate the possible immune effector mechanisms associated with the maladaptive response to *Mtb* infection in IFNγ^-/-^ T cell chimeric mice, we assessed gene expression in lung MDMs, the primary infected cell type in pulmonary TB. In FACS-sorted *Mtb*-mCherry infected and bystander MDMs at 25dpi, we noted prominent suppression of multiple hallmark M1 genes commonly associated with antimycobacterial responses, including *Nos2*, *IL12a*, and *IL12b,* and concurrent upregulation of a subset of hallmark alternative activation (M2) genes, including *Arg1*, *Chil3 (Ym1)*, *Mrc1 (CD206)*, *Fn1*, *Retnla (Fizz1)*, and *Ccl22* (**Fig. 4A**). Cytokine quantification in whole-lung lysates correlated the observed M2-related gene expression in IFNγ^-/-^ T cell chimera MDMs (**Fig. 4A**) with the presence of the canonical Th2 cytokines IL-4, IL-5, and IL-13, along with nearly complete absence of IFNγ (**Fig. 4B**). Confocal microscopy confirmed that, consistent with transcriptional data, expression of NOS2 was absent in lung MDMs of IFNγ^-/-^ T cell chimeric mice at 25dpi, while the M2 marker ARG1 was expressed on a much greater proportion of MDMs (**Fig. 4C**). We asked whether the type 1 interferon response, which has been associated with an ineffective and pathogenic response to *Mtb* infection(28–31) including neutrophil-associated pathology(32), may be responsible for the shutdown of type 2 interferon responses in IFNγ^-/-^T cell chimeras. However, transcription of both type 1 and type 2 interferon responses was suppressed in IFNγ^-/-^ T cell chimera mouse lung MDMs, suggesting that type 1 interferons do not play a major role in suppressing type 2 interferon-induced responses in these mice (**Fig. S4**).

**Figure 4:**
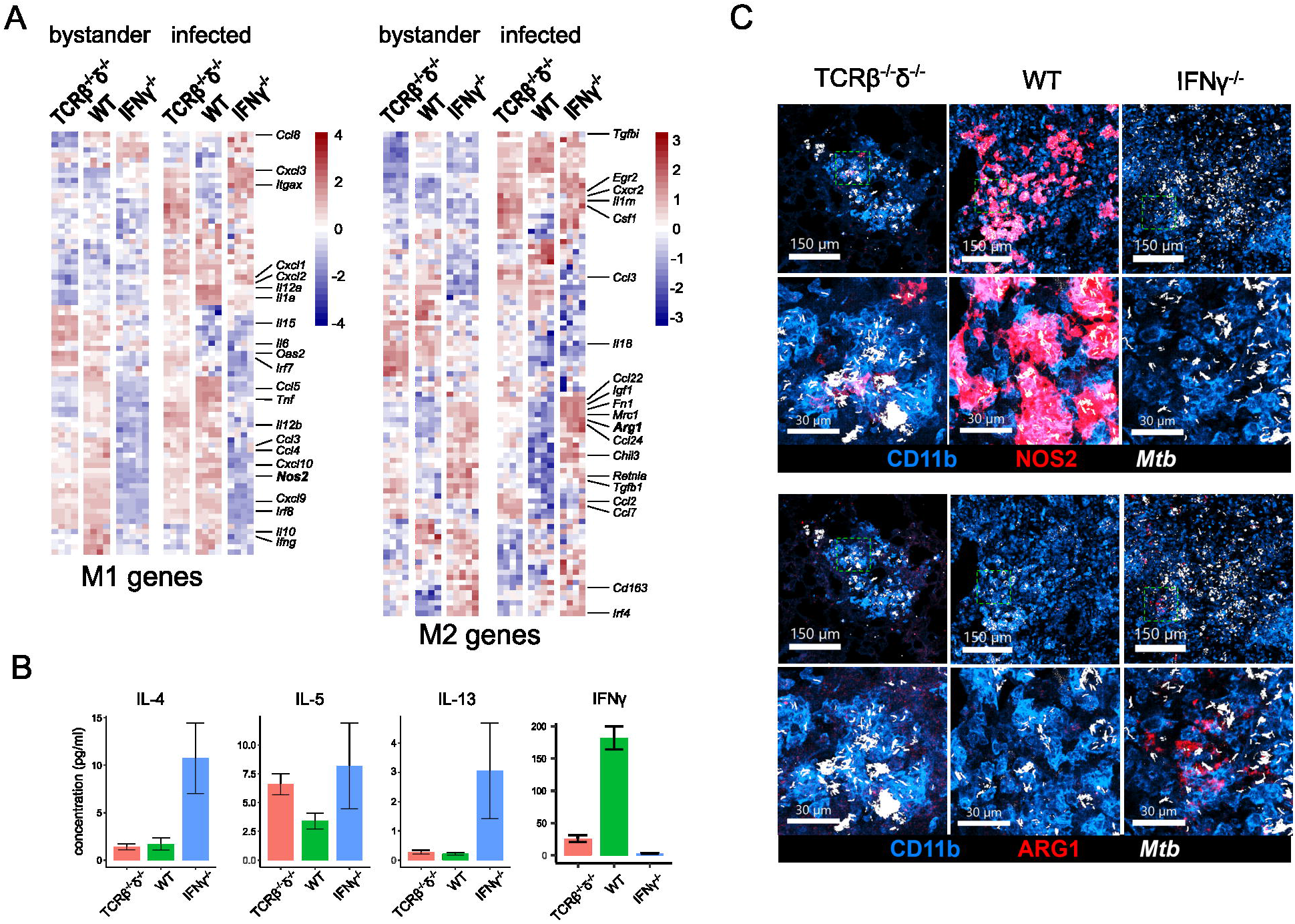
IFNγ^-/-^ T cells drive a Th2 cytokine milieu and alternative activation of monocyte-derived macrophages in T cell chimeric mice infected with *Mtb*. (A) Relative expression of classical (M1) and alternative (M2) genes in FACS-sorted bystander and *Mtb-*infected MDMs in lungs of T cell chimeric mice at 25dpi. (B) Concentration of Type 2 cytokines in lung lysates from *Mtb-*infected T cell chimeric mice at 25dpi, as measured by cytokine bead array. Results are representative of two independent experiments. (C) Concentration of IFNγ in lung lysates from *Mtb-*infected T cell chimeric mice at 25dpi, as measured by Luminex assay. Results are representative of two independent experiments. (D) Representative confocal microscopy demonstrating expression of iNOS and ARG1 in lungs of *Mtb-*infected T cell chimeric mice at 25dpi.

## DISCUSSION

Despite prior reports that T cell-derived IFNγ plays a minimal role in *Mtb* restriction in the C57BL/6 mouse model, our studies confirm it is indeed essential to control lung *Mtb* burden and disease in this setting. Furthermore, our results indicate a detrimental role for T cell signals unopposed by concomitant T cell-derived IFNγ in pulmonary *Mtb* infection. Unopposed IFNγ-independent T cell signaling correlates with clinical decline and recruitment of neutrophils and eosinophils to *Mtb* lung lesions. These effects did not involve a type 1 interferon response, but correlated with a Th2 cytokine signature and skewing of MDMs to an M2 phenotype. Pathology was worse in IFNγ^-/-^ compared to TCRβ^-/-^δ^-/-^ T cell chimeric mice despite no statistical difference in lung *Mtb* CFU burden, suggesting that T cell-derived IFNγ regulates tolerance of the host to manifestations of disease from *Mtb* infection, such as tissue damage and weight loss. An effective vaccine strategy against TB disease will likely require both IFNγ-dependent and independent T cell effects, and that – as is frequently the case for immune responses in vivo – a balance between these two arms of the T cell response is required for effective protection while minimizing immunopathology.

Pulmonary lesions in TB disease-susceptible IFNγ^-/-^ T cell chimeric mice were marked by neutrophil and eosinophil infiltration. Whether the recruitment of these cell types is a major driver of lung pathology and clinical deterioration remains to be determined. While neutrophils may play a host-protective role in mycobacterial clearance early in infection through phagocytosis and ROS production, adverse outcomes in Mtb infection are usually associated with a dysregulated neutrophil response that plays a major role in driving detrimental pathology(33). Prior studies have also shown that IFNγR^-/-^ neutrophils accumulate in the lungs of *Mtb-*infected WT/IFNγR^-/-^ mixed bone marrow chimeric mice, and that these mice exhibit accelerated weight loss(15), consistent with a direct role for IFNγ in suppressing harmful neutrophil accumulation in the lung during *Mtb* infection. Our results build on this model by suggesting that T cells may provide an essential source of IFNγ that inhibits the pathogenic effects of neutrophils. Whether the abundant eosinophils we observed in pulmonary TB lesions of IFNγ^-/-^ T cell chimeric mice also drive detrimental pathology, or a potentially mediate a host-beneficial response to severe disease, deserves further study. Prior work using two independent genetic models of global eosinophil deficiency demonstrated a role for these cells in restriction of mycobacterial burden(34). However, the contribution of eosinophils to the host-pathogen balance may be context-specific.

Lung lysates of *Mtb*-infected IFNγ^-/-^ T cell chimeric mice were marked by a significant Th2 cytokine profile, with abundant IL-4, IL-5, and IL-13. In prior murine studies, depletion of IL-4 led to improved control of *Mtb* burden in BALB/c mice (35). In C57BL/6 mice, known to have a strong Th1-skewed response to *Mtb* infection, pulmonary *Mtb* burden was not affected by global deficiency in IL-4 or IL-13(36). However, over-expression of IL13 in C57BL/6 mice led to formation of necrotizing granulomas(37) in an IL-4Rα-dependent manner(38). Further, alternatively activated M2 macrophages expressing ARG1, associated with the presence of Th2 cytokines, were abundant in this model(37). Accordingly, we observed suppression of the Th1-driven NOS2 and induction of the Th2-driven M2 marker ARG1 in macrophages in *Mtb* lesions of IFNγ^-/-^ T cell chimeric mice. Consistent with these findings, Th2 responses have been shown to be responsible for more severe pulmonary inflammation and TB disease in mice exposed to *Schistosoma mansoni* parasites or antigen than in untreated mice across a range of murine genotypes (39). Together, this evidence again suggests a context-specific effect that may depend on factors including genetic background, helminth coinfection, and environment.

There is also mounting evidence from human clinical and experimental studies supporting a detrimental role for Th2 cytokine signaling in TB pathology(40). A study of 1971 HIV-negative patients with sputum culture-positive pulmonary TB in Ghana revealed that a variant of IL4-Rα associated with increased signal transduction was associated with increased cavity size(38). In addition, a significantly higher IL-4/IFNγ ratio was observed in stimulated lung lymphocytes from bronchoalveolar lavage (BAL) in patients with miliary(41) or cavitary(42) rather than pleural(41) or non-cavitary(42) TB. The same pattern was seen among peripheral blood lymphocytes(43), though the proportion of circulating IL-4-expressing T cells was much smaller than in BAL when measured concurrently in the same patients(41), implying an important role for tissue-specific responses. Indeed, in-situ hybridization using resected lung tissue from patients with severe pulmonary TB demonstrated IFNγ and IL-4 mRNA-producing cells within the same granulomas, as well as co-existence of these mixed granulomas alongside granulomas expressing only IFNγ within the same patient(44); each lesion may therefore represent a unique cytokine micro-environment. Thus, while Th2-associated comorbidities such as helminth infection or allergy/atopy have not consistently correlated with severity of TB disease(45–47), studies have repeatedly demonstrated an association between the ratio of Th2 vs Th1 cells and severity of *Mtb* infection outcomes. It has been suggested that a vaccine that would protect against *Mtb* in areas where both TB and helminthic infections are endemic should both support the Th1 response and block the Th2 response(40, 48). Our findings correlating Th2 cytokines and M2 macrophage responses with severe TB disease support this framework.

While T cell-derived IFNγ is most often thought of in terms of its effect on macrophages, it is not known which cellular target is primarily responsible for the differences in *Mtb* infection outcome between WT and IFNγ^-/-^ T cell chimeric mice. T cell-derived IFNγ can be sensed by T cells themselves, and plays a role in subsequent Th1 polarization(49). Furthermore, effects of IFNγ on *Mtb* immunity have been correlated with induction of gene expression in epithelial cells(50), and by shaping T cell compartments by inducing apoptosis of activated CD4+ T cells(51). Future studies to characterize cell compartments, apoptosis, and activation status in lungs of *Mtb*-infected IFNγ^-/-^ T cell chimeric mice will help address this question.

In a prior study, depletion of CD4+ T cells led to worsening *Mtb* infection outcomes in mice despite relatively preserved levels of total IFNγ in the lung(20). Notably, in our studies, production of lung IFNγ and expression of NOS2 by lung MDMs was even lower in IFNγ^-/-^T cell chimeric mice than in T cell chimeras that lacked T cells completely. This suggests that there is IFNγ production by cell types other than T cells, such as NK or monocytic cells, which may expand in a compensatory manner when T cells are absent – but these cells may be suppressed or unable to meaningfully increase IFNγ production when T cells lack IFNγ. It is plausible that this striking difference may be due to the fact that IFNγ^-/-^ T cell chimeric mice had never possessed IFNγ-competent T cells, while mice in the CD4+ T cell depletion studies were previously exposed to T cell-derived IFNγ, and may have therefore achieved a baseline Th1 T cell driven response or tonic state that could support the production of IFNγ by other cell types when needed.

Prior studies showed that IFNγ produced by adoptively transferred Th1-polarized, *Mtb* antigen-specific CD4+ T cells is dispensable to control of pulmonary *Mtb* burden in WT host mice(6) when transferred at a dose of 1×10^7^ cells/host, and partially dispensable at a dose of 1×10^6^ cells/host. Together with our results, these data indicate that high numbers of *Mtb* antigen-specific, Th1-polarized IFNγ^-/-^CD4+ T cells likely amplify the importance of IFNγ-independent T cell effects and overcome a requirement for T cell-derived IFNγ to reduce *Mtb* burden in the lung, while dependence on T cell-derived IFNγ is unmasked at more physiologic numbers of antigen-specific T cells that are more closely aligned with an expected vaccine response.

One factor that limits the interpretation of our studies is that C57BL/6 mice, the strain used in our work, is known to elicit a Th1-skewed response, whereas other genetic backgrounds may be less dependent on IFNγ for protection against TB (52, 53). Furthermore, Th1-driven mechanisms in C57BL/6 mice may differ from those in humans. While the role of IFNγ-induced NOS2 in *Mtb* restriction is well-established in mice, *Mtb*-infected human peripheral blood-derived monocytes produce only small amounts of nitric oxide in response to IFNγ signaling in vitro(54). Whether NOS2 is induced at sites of *Mtb* infection in human lungs is controversial, with studies showing different results(55–57). Nevertheless, the most frequently cited manuscripts arguing for a minimal role of T cell-derived IFNγ in pulmonary immunity against TB are studies in C57BL/6 mice, which are inconsistent with our results.

Our study was performed in unvaccinated mice, and it remains possible that vaccination could boost other mechanisms of immunity independently of T cell-derived IFNγ. For example, though prior studies have definitively proven the requirement for IFNγ in control of *Mtb*, IFNγR^-/-^ mice vaccinated with BCG still have a survival advantage over unvaccinated IFNγR^-/-^ mice(17); while T cell-independent effects such as trained immunity could be responsible, IFNγ-independent T cell activity may also play a role.

Nevertheless, our results indicate that T cell-derived IFNγ can be critical for immunity within the *Mtb*-infected lung and vaccination or host-directed therapy strategies that restore or augment the ability of *Mtb*-specific T cells to produce IFNγ should continue to be explored.

## ACKNOWLEDGMENTS

We thank Daniel Kim, Lindsay Engels, Kaitlin Durga, and the SCRI Animal Care staff for technical assistance, Alan Diercks for RNA-seq data alignment, SCRI Research Scientific Computing for HPC resources, and other members of the Urdahl Lab for helpful discussions. This study was supported by NIH grants U19AI135976 (K.B.U.), 75N93019C00070 (K.B.U.), and T32AI007044 (K.M.), the Firland Foundation 20230026C (K.M.), the American Lung Association CAALA2023 (K.M.), and the NIH-funded Seattle TB Research Advancement Center (SEATRAC) 1P30AI168034-01 (K.M.). The sponsors had no role in the design, conduct, analysis, or interpretation of the study, nor in the preparation, review, or approval of the manuscript.

## Notes

### Competing Interest Statement

The authors have declared no competing interest.

