## Supplemental Figures for "Re-appraising the role of T-cell derived interferon gamma in restriction of *Mycobacterium tuberculosis* in the murine lung"

Fig. S1

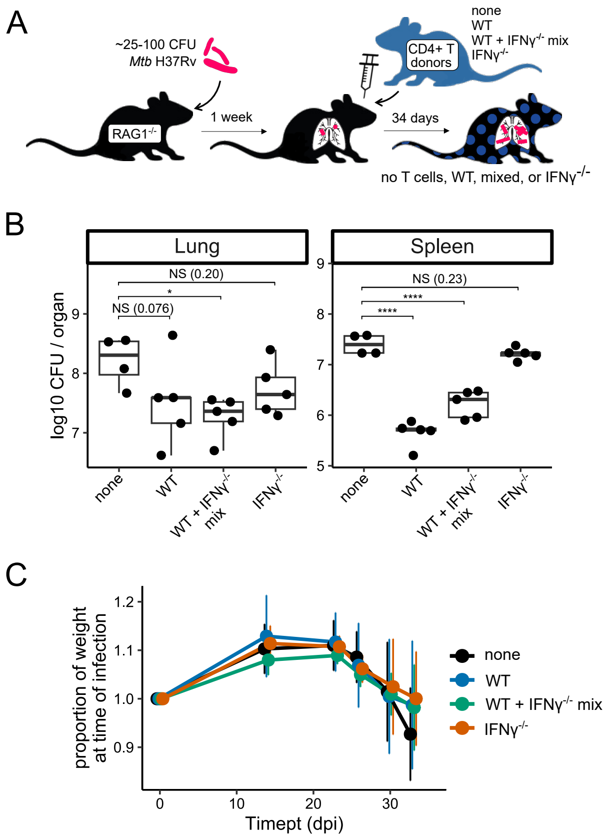


**Figure S1: Adoptively transferred IFNγ$^{-/-}$ CD4+ T cells do not reduce *Mtb* burden, but do not affect tolerance to infection relative to WT T cells in RAG1$^{-/-}$ mice.**

(A) Schematic of adoptive transfer of zero or 3*10$^{6}$ WT, 50/50% mixed, or IFNγ$^{-/-}$ CD4+ T cells to RAG1-deficient host mice after infection with aerosolized *Mtb*.

(B) Bacterial burden in lungs and spleens of *Mtb*-infected adoptive transfer mice 34dpi. Results are representative of two independent experiments. Statistical significance was determined by Tukey's range test. *$p \leq 0.05$, **$p \leq 0.01$, ***$p \leq 0.001$, ****$p \leq 0.0001$.

(C) Weight trends of *Mtb*-infected adoptive transfer mice through 34dpi.

Fig. S2

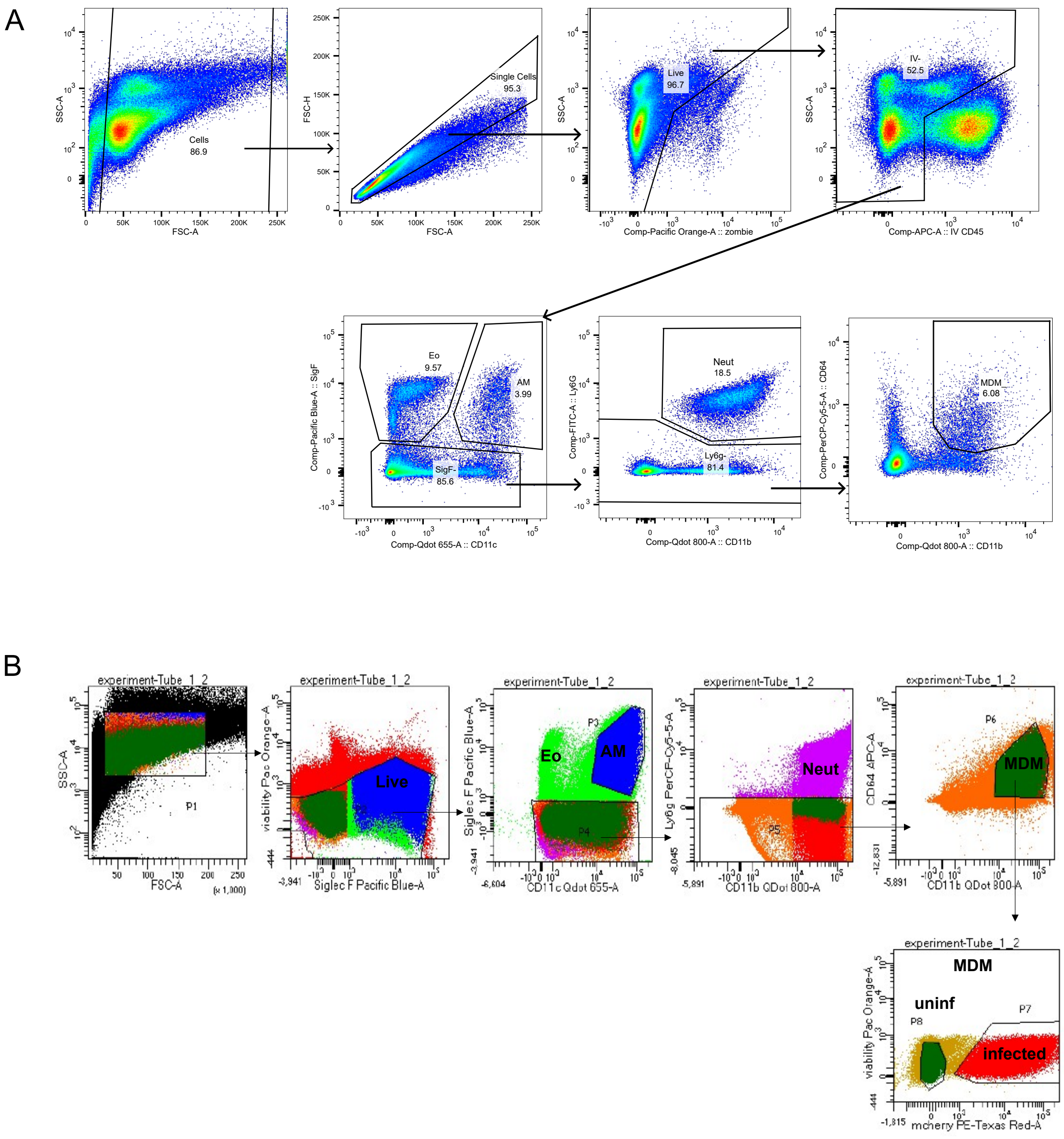


**Figure S2: Flow cytometry gating to identify myeloid subsets in lungs of *Mtb*-infected mice.** T cell chimeric mice were infected with *Mtb* H37Rv-mCherry, IV-labeled, and harvested at 25dpi. Single cell suspensions from the right lung were stained with antibodies targeting myeloid cell subsets and gated as shown for flow cytometric analysis (A) and cell sorting prior to RNA-seq (B).

Fig. S3

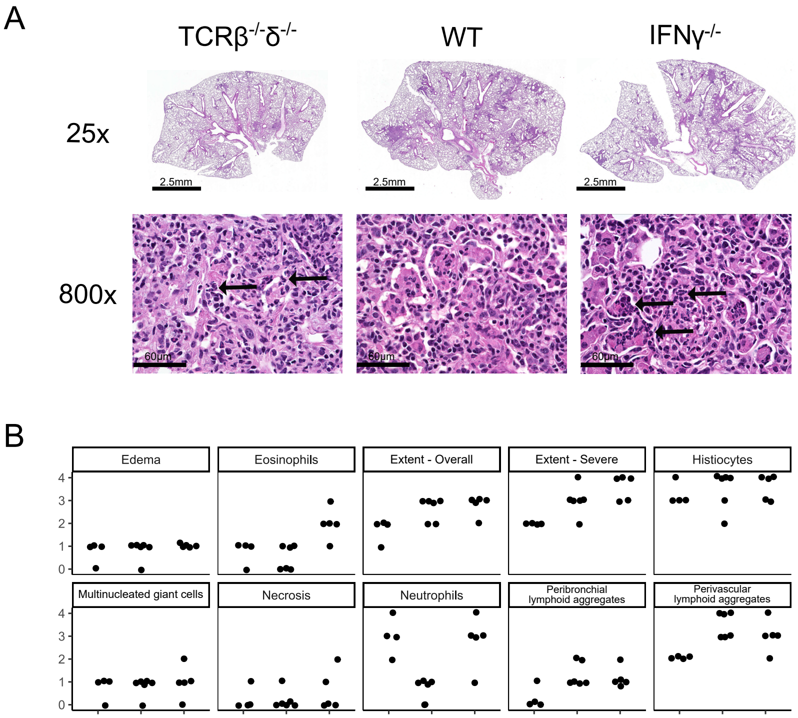


**Figure S3: Histopathologic scoring of representative sections of H&E stained lung sections from T cell chimeric mice infected with *Mtb.***
Histopathologic features assessed in representative sections of fixed and hematoxylin-eosin (H&E) stained lung of T cell chimeric mice at 25dpi. Sections were scored by a pathologist blinded to T cell chimera genotype of each sample. Each dot represents one mouse. MNGC: multinucleated giant cells. PB: peribronchial. PV: perivascular. LA: lymphocyte aggregates.

### Fig. S4

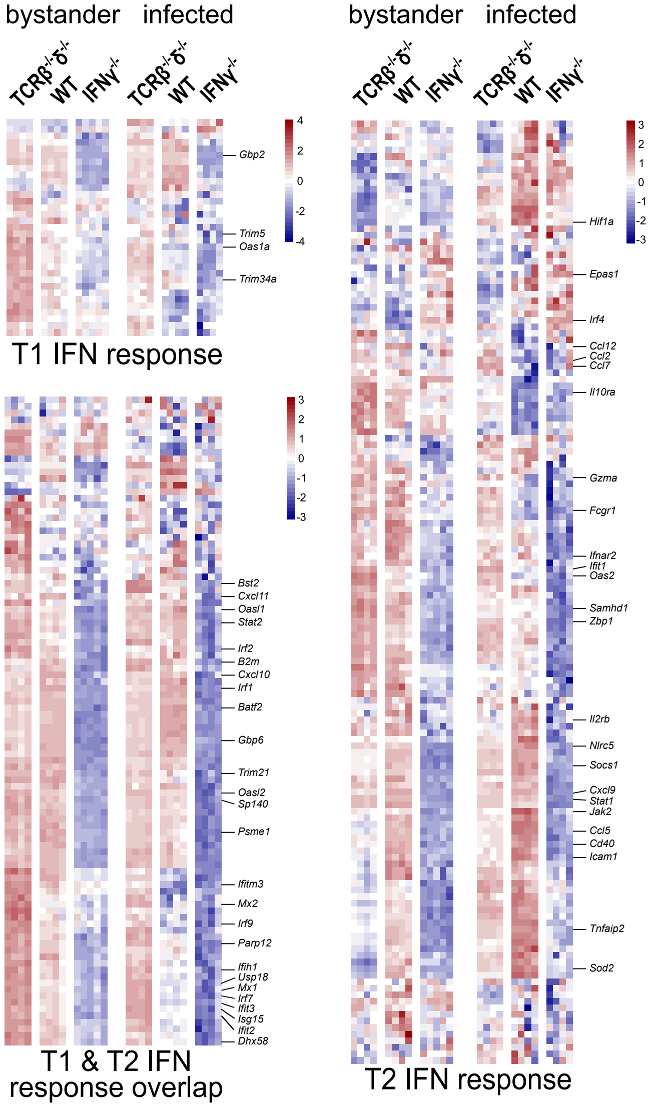


**Figure S4: IFNγ$^{-/-}$ T cells suppress both Type 1 and Type 2 interferon responses in monocyte-derived macrophages in T cell chimeric mice infected with *Mtb*.** Row-normalized relative expression of Type 1 and 2 interferon-induced genes in FACS-sorted bystander and infected MDMs in lungs of T cell chimeric mice at 25dpi.
